# Rapid generation of circulating and mucosal decoy ACE2 using mRNA nanotherapeutics for the potential treatment of SARS-CoV-2

**DOI:** 10.1101/2020.07.24.205583

**Authors:** Jeonghwan Kim, Anindit Mukherjee, Dylan Nelson, Antony Jozic, Gaurav Sahay

## Abstract

Severe acute respiratory syndrome coronavirus 2 (SARS-CoV-2) enters through the airways and infects the lungs, causing lethal pulmonary damage in vulnerable patients. This virus contains spike proteins on its envelope that binds to human angiotensin-converting enzyme 2 (hACE2) expressed on the surface of airway cells, enabling entry of the virus for causing infection^1,2^. In severe cases, the virus enters the circulatory system, contributing to multiorgan failure. Soluble form of hACE2 binds to SARS-CoV-2 spike protein and prevents viral entry into target cells^3^. Moreover, soluble recombinant ACE2 ameliorates lung injury^4^ but its short half-life limits its therapeutic utility^5^. Here, we engineered synthetic mRNA to encode a soluble form of hACE2 (hsACE2) to prevent viral infection. Novel lipid nanoparticles (LNPs) were used to package mRNA and transfect mammalian cells for enhanced production of secreted proteins. Intravenously administered LNP led to hepatic delivery of the mRNA. This elicited secretion of hsACE2 into the blood circulation within 2 h, and levels of circulating hsACE2 peaked at 6 h and gradually decreased over several days. Since the primary site of entry and pathogenesis for SARS-CoV-2 is the lungs, we instilled LNPs into the lungs and were able to detect hsACE2 in the bronchoalveolar lavage fluid within 24 h and lasted for 48 h. Through co-immunoprecipitation, we found that mRNA-generated hsACE2 was able to bind with the receptor binding domain of the SARS-CoV-2 spike protein. Furthermore, hsACE2 was able to strongly inhibit (over 90%) SARS-CoV-2 pseudovirus infection. Our proof of principle study shows that mRNA-based nanotherapeutics can be potentially deployed for pulmonary and extrapulmonary neutralization of SARS-CoV-2 and open new treatment opportunities for COVID-19.

## MAIN

More than 15 million cases with over 600,000 deaths have resulted from the ongoing coronavirus disease 2019 (COVID-19)^6^. SARS-CoV-2, the pathogen of COVID-19, is a β-coronavirus that primarily enters through the airways and lungs. The envelop of SARS-CoV-2 is decorated with homotrimeric spike (S) proteins that bind to the human angiotensin-converting enzyme 2 (hACE2) receptor expressed on the cell surface^1,2^. The S protein is composed of S1 and S2 subunits responsible for viral attachment and fusion^7^, respectively. Binding between the receptor-binding domain (RBD), which is located within the S1 subunit, and hACE2 triggers a cascade that accelerates cellular entry and viral membrane fusion. hACE2 is expressed in the lungs, heart, kidney, and intestine^8^. The inherent function of hACE2 is a key enzyme that participates in the Renin Angiotensin Aldosterone System (RAAS) responsible to maintain blood pressure. hACE2 is a carboxypeptidase that converts Angiotensin 1 to Angiotensin (1-9) or Angiotensin II to Angiotensin (1-7), both of which are vasodilators with cardioprotective effects through regulation of blood pressure^9^. SARS-CoV-2 interacts with hACE2 to enter and infect human airway epithelial cells, causing cytotoxic responses. It also can develop pneumonia and cytokine storm, leading to Acute Respiratory Distress Syndrome (ARDS) in severe cases^10–12^. Once the virus infiltrates systemic circulation, it can dysregulate RAAS and immune system, cause endothelial cell damage, possibly targets other tissues that express hACE2, and overall cause a multiorgan failure^13^. hACE2 consists of three segments: an extracellular segment that contains the peptidase domain where the RBD binds to, a transmembrane segment, and an intracellular segment^2^. hACE2 can be cleaved by peptidases at the neck region of the extracellular segment, releasing a soluble form of hACE2 (hsACE2) which is enzymatically active^14,15^. Since the RBD of SARS-CoV-2 binds to the extracellular domain of hACE2, hsACE2 protein can prevent the viral infection through competitive inhibition^14,15^. Monteil et al. recently demonstrated that recombinant hsACE2 can inhibit the infection of SARS-CoV-2 in cultured kidney organoids^3^. However, a relatively short half-life of the recombinant hsACE2 in the bloodstream would necessitate repeated administrations to ensure long-term circulation of the protein for days after exposure to SARS-CoV-2^16^. Chimeric hsACE2 that contains a fused Fc region of human IgG, reduced viral infectivity *in vitro*; however, its pharmacokinetic benefit remains to be determined^17^. A short half-life of soluble ACE2 (< 2 h in mouse) severely limits its time window of action^5^ and extended residence of hsACE2 is desirable to mitigate SARS-CoV-2 mediated RAAS activation and hence to reduce inflammation-related injury of organs. To overcome these challenges, we use LNPs to deliver *in-vitro-transcribed* messenger RNA (IVT mRNA) for rapid expression of hsACE2 **(Fig. 1a-c**). This strategy may allow for rapid clearance of the captured virus while maintaining hsACE2 levels that can surveil circulation, clear the virus, and rescue the disrupted RAAS system. mRNA-based therapy has been attractive since the process of mRNA synthesis became fast, affordable, and scalable^18^, and these advantages have enabled the fast development of mRNA vaccines for COVID-19^19,20^. Moreover, LNP-delivered mRNA provides a transient yet high expression of protein with proper folding and post-translational modifications, but without risk of insertional mutagenesis as is associated with viral-based gene therapy^21^. Unlike viral vectors, this platform technology can be repeatedly administered to sustain protein production until the infection subsides and cease of the treatment allows for clearance of hsACE2 within days, mitigating any off-target effects. We hypothesized that expression of hsACE2 will prevent SARS-CoV-2 from binding to cell surface receptors and block its entry **(Fig. 1a, b)**. We designed IVT mRNA to encode the 1-740 amino acid sequence of hACE2 with a cleavable V5-epitope tag at the C-terminus **(Fig. 1c and Extended Data Fig. 1)**.

**Fig. 1.**
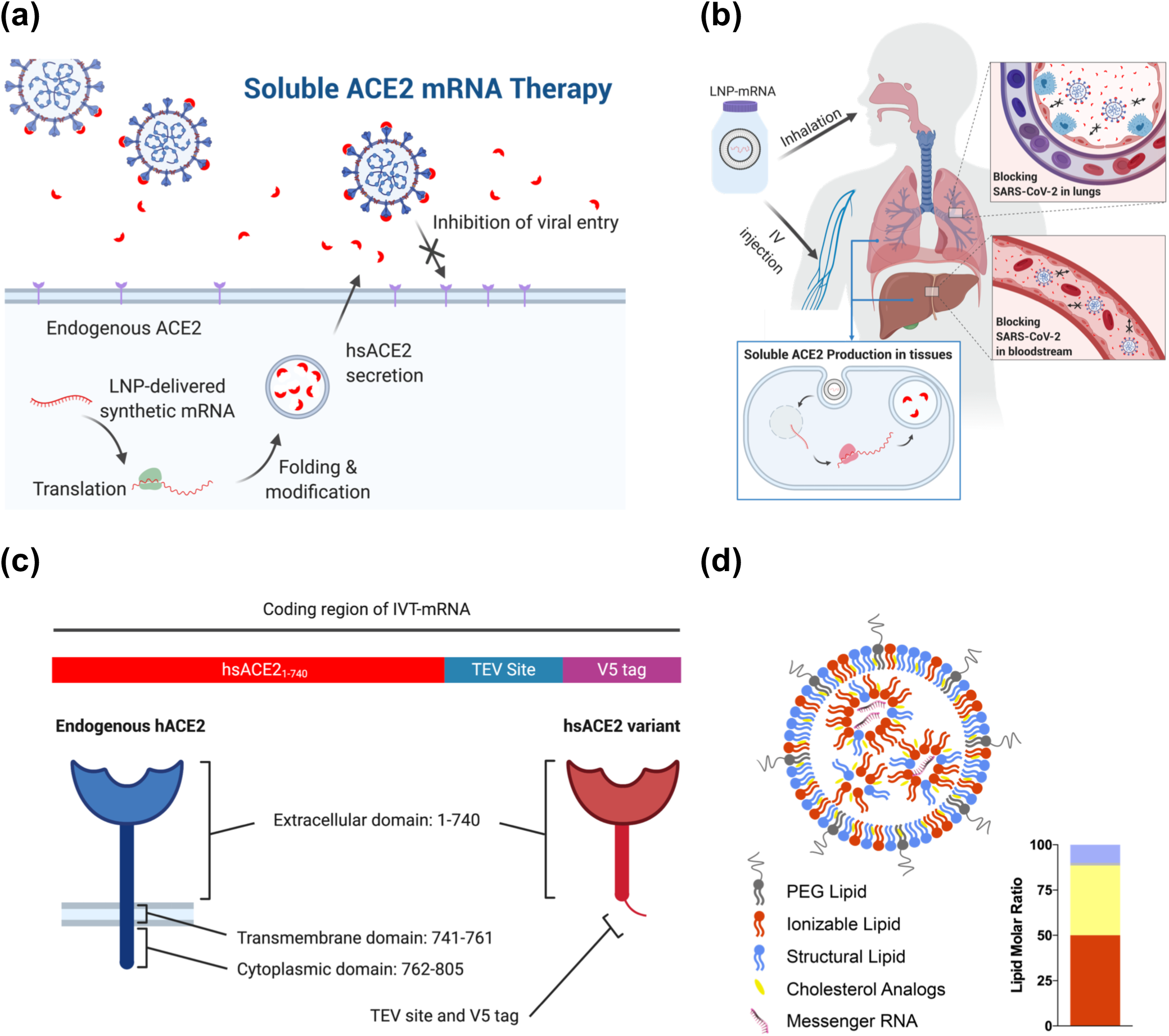
Design of mRNA-based nanotherapeutic to treat SARS-CoV-2 infection. (a) Rationale for soluble ACE2 mRNA therapeutic in treating SARS-CoV-2 infection. LNP-delivered synthetic mRNA generates human soluble ACE2 (hsACE2) protein that is secreted into the extracellular compartment where it binds receptor binding domain of the spike protein of the SARS-CoV-2 and prevents viral entry. (b) Potential routes of administration for soluble ACE2 mRNA therapeutic. Intravenous administration of the LNPs encapsulating *hsACE2* mRNA leads to the production of circulating hsACE2 protein from the liver. Inhalation of the LNPs leads to production of mucosal hsACE2 protein in the lungs. (c) Schematic of *in-vitro*-transcribed (IVT) mRNA encoding hsACE2 variant protein (red), TEV site (blue), and V5 tag (purple) (top). Endogenous hACE2 consists of three domains: extracellular, transmembrane, and cytoplasmic domains (bottom left). The hsACE2 variant lacks transmembrane and cytoplasmic domains of the endogenous hACE2 protein, and it contains TEV site and V5-epitope tag at C-terminus (bottom right). (d) Schematic representation of LNP encapsulating mRNA. Inset, molar ratio of each lipid in the LNP. Orange; ionizable lipid, Yellow; cholesterol analogs, Blue; structural lipid, Gray; PEG lipid.

To confirm whether the designed IVT mRNA produces hsACE2 protein after transfection, 293T cells were transfected with *hsACE2* mRNA using lipofectamine 3000. We detected the presence of hsACE2 protein in cell-free conditioned media and cell lysates from *hsACE2* mRNA transfected cells by western blot, but not in PBS-treated controls **(Extended Data Fig. 2a, b)**. Efficient intracellular delivery of mRNA, especially *in vivo*, requires a potent delivery vector. Conventional LNPs are composed of four lipids: (1) ionizable lipid, (2) PEG lipid, (3) cholesterol, and (4) structural lipid **(Fig. 1d)**. Recently, we discovered a simple substitution of cholesterol to β-sitosterol within LNP formulations boosts intracellular delivery of mRNA^22,23^. We compared the enhanced LNP formulation (eLNP: containing β-sitosterol) with the conventional LNP (containing cholesterol) as the delivery vector for *hsACE2* mRNA. Both LNP and eLNP encapsulating *hsACE2* mRNA (LNP or eLNP/hsACE2) exhibited comparable characteristics in terms of hydrodynamic sizes (≈ 80 nm), polydispersity (PdI ≈ 0.08), and RNA encapsulation (above 98%) **(Extended Data Fig. 2c, d)**. We found that eLNP encapsulating firefly luciferase (*Fluc*) mRNA generated significantly higher luciferase expressions than LNP (p<0.01) with a dose-dependent manner in 293T cells without decreasing cell viability, indicating improved transfection efficiency **(Extended Data Fig. 3a, b)**. In *hsACE2* mRNA delivery, eLNP/hsACE2 elicited substantially greater production of hsACE2 protein compared to LNP/hsACE2 in cell-free conditioned media from 293T culture. In the cell-free conditioned media from 293T cells treated with LNP/Fluc and eLNP/Fluc, no expression of hsACE2 was detected in western blot **(Extended Data Fig. 3c)**. The improved hsACE2 expression by eLNP was confirmed again in the analysis of cell lysates (5-fold higher expression, p < 0.0001) **(Extended Data Fig. 3d, e)**. Additionally, the production of hsACE2 protein was dependent on the mRNA dose given **(Fig. 2a)**.

**Fig. 2.**
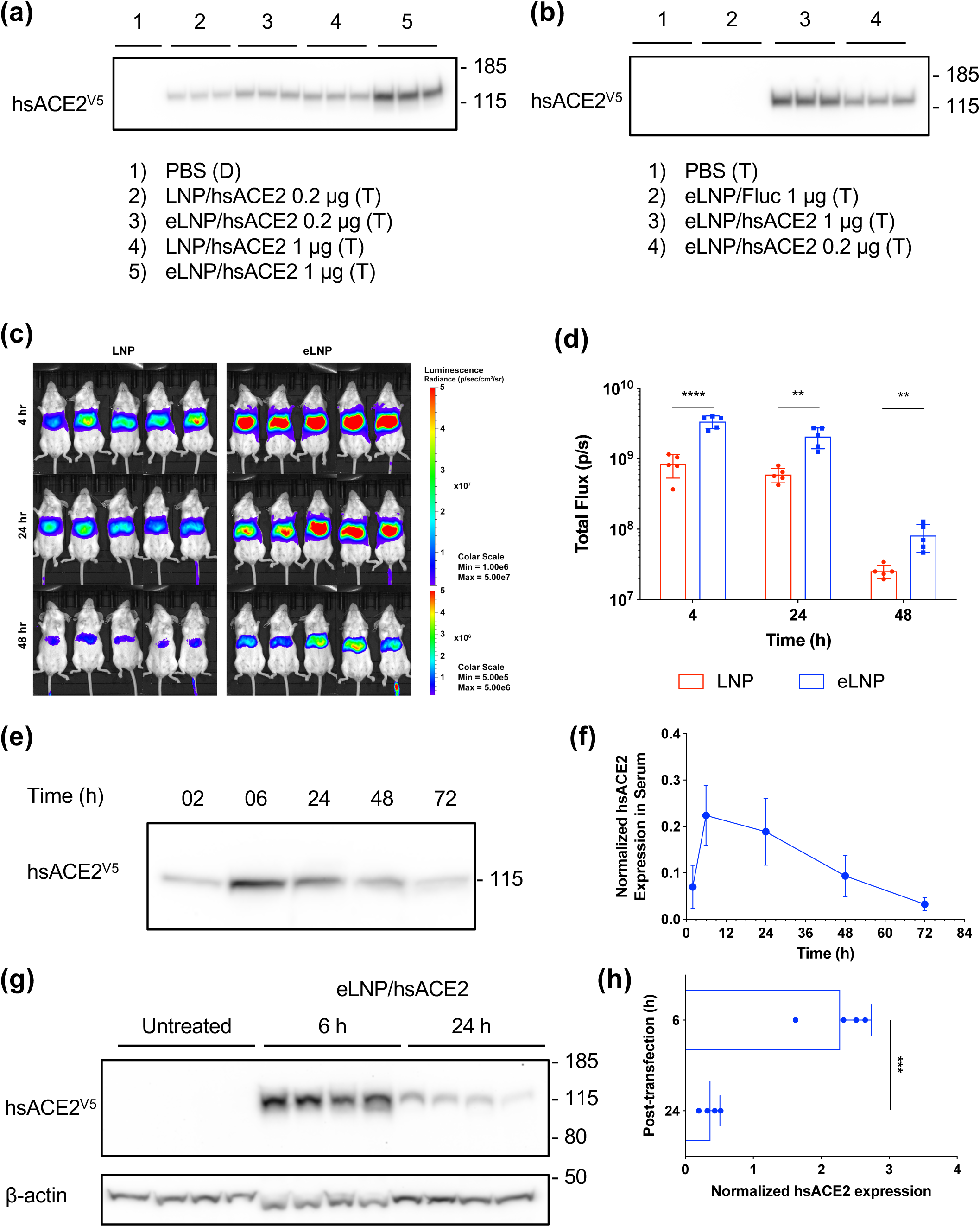
*hsACE2* mRNA transfection result in rapid production of the circulatory hsACE2 protein. (a, b) Western blot of cell-free conditioned media from (a) 293T and (b) Hep G2 cell culture after mRNA transfection using various LNPs. Treatment and mRNA dose are described under each blot (Duplicate (D): n=2, Triplicate (T): n=3). (c) *In vivo* bioluminescent images of BALB/c mice after treatment of 0.05 mg/kg mRNA delivered through IV injection of LNP/Fluc or eLNP/Fluc (n = 5). (d) Quantification of bioluminescent signals from IVIS images. Region of interest kept constant in all images (n = 5). (e) A representative image of western blot with mouse sera collected with predetermined time intervals after IV injection of 0.15 mg/kg eLNP/hsACE2. (f) Densitometric quantitation of temporal levels of the circulatory hsACE2 protein in mouse sera after IV injection of eLNP/hsACE2 (n = 5). Expression of hsACE2 protein was normalized to the total amount of protein in each lane. All data were expressed as the mean ± S.D. (g) Western blot of liver homogenates collected from BALB/c mice after IV injection of 0.15 mg/kg eLNP/hsACE2. (h) Expression of hsACE2 protein in mouse liver homogenates after IV injection of eLNP/hsACE2 (n = 4). Densitometric analysis of hsACE2 protein expression normalized to β-actin levels. Statistical analysis was performed using Student’s *t* test. \*\**p* < 0.01, \*\*\**p* < 0.001, \*\*\*\**p* < 0.0001. All data were expressed as the mean ± S.D.

Accumulating evidence suggests that invasion of SARS-CoV-2 in ACE2-expressing airway epithelial cells is followed by infection of endothelial cells, leading to endotheliitis^24,25^. Vascular leakage caused by damaged endothelial cells provides the virus with a putative gateway to the circulatory system and other ACE2-expressing organs^25^. Therefore, blockade of influx of the virus from the blood circulation to peripheral organs is likely to prevent multisystem organ failure. For these reasons, we evaluated *in vivo* delivery of eLNP/hsACE2 for production, secretion, and blood circulation of hsACE2 protein. Intravenously administered LNPs are typically destined to transfect hepatocytes owing to the interaction between LNP and apolipoprotein E^26^. Thus, we expected the liver will serve as a factory for protein production upon *hsACE2* mRNA transfection as shown in other studies for secretory proteins^27,28^. To demonstrate this *in vitro*, the human liver cell line, Hep G2, was transfected with LNPs. eLNP/Fluc yielded more luciferase expression than LNP/Fluc in Hep G2 cells **(Extended Data Fig. 4a, b)**. As was the case with 293T cells, hsACE2 protein was found within the cell-free conditioned media and lysates that were harvested from the transfected Hep G2 cells in a dose-dependent manner **(Fig. 2b and Extended Data Fig. 4c, d)**. Interestingly, in the Hep G2 cell lysates, two discrete hsACE2 bands were observed. We assume the band at approximately 125 kDa represented the fully-glycosylated form and the band at 100 kDa represented the pre- or partially-glycosylated form **(Extended Data Fig. 4c)**. However, only the fully-glycosylated form was detected in the cell-free media **(Fig. 2b)**, suggesting the glycosylation of hsACE2 protein is prior to secretion. We then examined whether the eLNP lead to improved protein production *in vivo*. It was shown that eLNP/Fluc induced strong bioluminescent signals in the livers of BALB/c mice after intravenous injection of LNPs at 4 hours post-injection, which decreased with time (**Fig. 2c)**. eLNP/Fluc exhibited 3-fold increase in luciferase expression as compared to LNP/Fluc at all time points (4-48 h) **(Fig. 2d)**. Based on these results, we used eLNP as an optimized formulation for delivering mRNA in the remaining studies. We injected eLNP/hsACE2 in BALB/c mice and collected mouse sera up to 72 h post-administration with predetermined time intervals. Notably, hsACE2 appeared in the mouse sera as early as 2 h post-injection **(Fig. 2e and Extended Data Fig. 5b, c)**. This rapid generation of hsACE2 from the liver will be useful to neutralize SARS-CoV-2 promptly at a stage of systemic spread. We found that hsACE2 was detected at the highest level at 6 h post-injection and gradually declined afterwards. We could detect circulating hsACE2 even 72 h after a single injection **(Fig. 2e, f and Extended Data Fig. 5a)**. The expression of hsACE2 protein in liver homogenates was time-dependent, showing a greater expression of the protein after 6 h than 24 h post-administration (p < 0.001) **(Fig. 2g, h)**. Unlike the cell lysates of Hep G2, the mouse liver homogenates showed a single band at approximately 125 kDa. After 7 days, hsACE2 was mostly eliminated from the blood circulation (data not shown).

Airway and lungs are the first target organs where the virus attacks and are highly vulnerable organs due to high levels of hACE2 expression^14^. Having hsACE2 protein as a decoy on the airway epithelium could mitigate viral infection at early stages of disease progression. Therefore, we explored the ability of LNPs to produce mucosal hsACE2. Consistent with the previous results, eLNP/Fluc exerted significantly greater levels of transfection than LNP/Fluc in Calu-3, a human lung epithelial cell line **(Fig. 3a, Extended Data Fig. 6a)**. Similarly, eLNP/hsACE2 showed substantially higher expression of hsACE2 protein than LNP/hsACE2 in western blot **(Fig. 3b and Extended Data Fig. 6c, d)**. To locally deliver LNPs to the mouse lungs, we used intratracheal instillation as the route of administration. Intratracheally administered eLNP/Fluc transfected the lungs of BALB/c mouse, and the luciferase expression was detected exclusively in the lungs **(Fig. 4c, d)**. To evaluate the secretion of hsACE2 to the lung mucosa, bronchoalveolar lavage fluid (BALF) samples were collected at 24 h and 48 h post-administration of eLNP/hACE2, followed by pull-down of hsACE2 protein using anti-V5 antibody. We confirmed the presence of mucosal hsACE2 in the collected BALF samples by western blot **(Fig. 3e)**. These data represent that lung transfection with eLNP/hsACE2 resulted in the secretion of hsACE2 to the airway mucus.

**Fig. 3.**
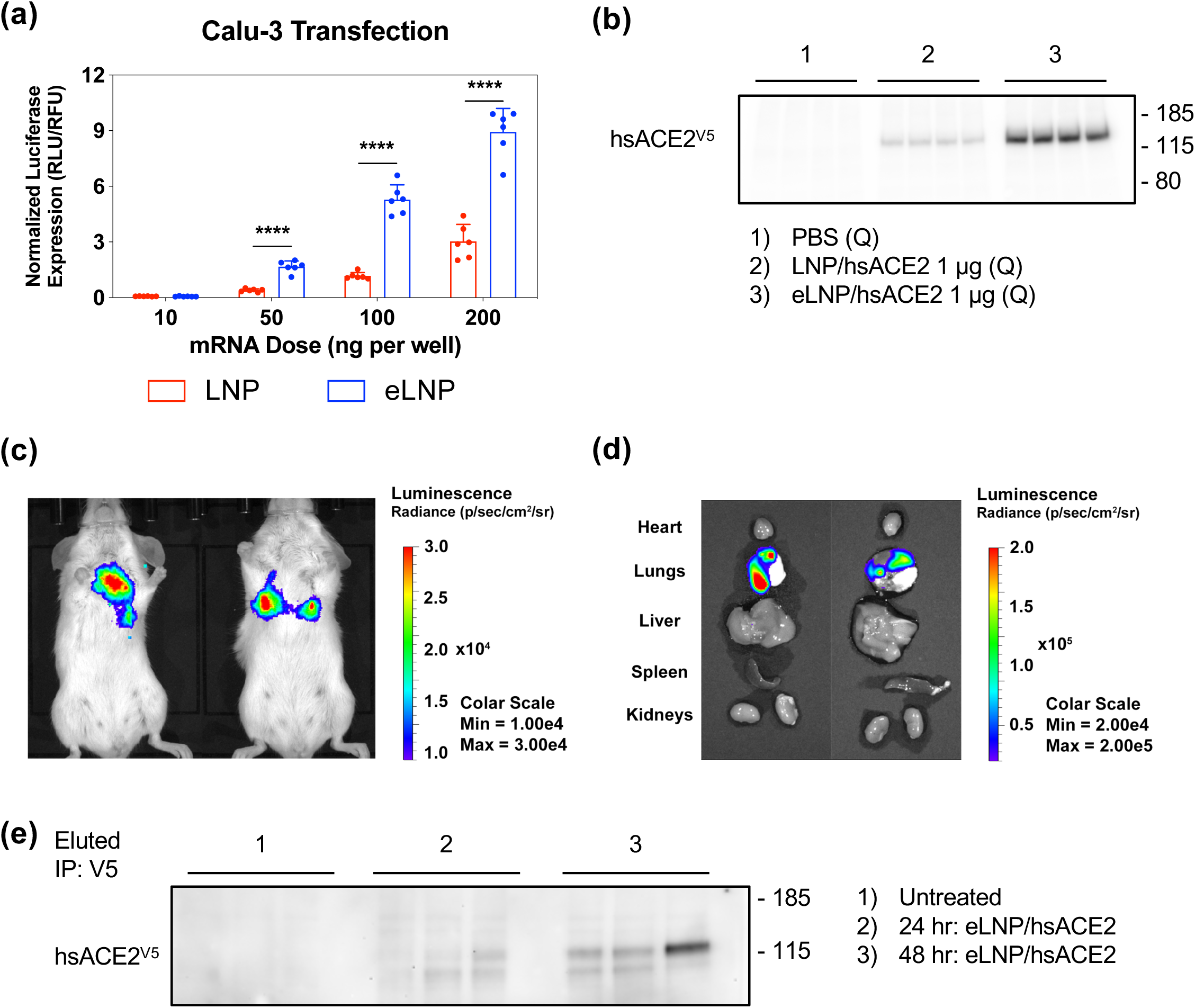
LNP-delivered *hsACE2* mRNA to the lungs resulted in production of mucosal hsACE2 protein. (a) *In vitro* transfection of Calu-3 by 48-hour incubation of LNP/Fluc or eLNP/Fluc at various mRNA doses (n = 6). Statistical analysis was performed using Student’s *t* test. \*\*\*\**p* < 0.0001. All data were expressed as the mean ± S.D. (b) Western blot of cell-free media of Calu-3 cell culture after *hsACE2* mRNA transfection using LNPs. Treatment and mRNA dose are described under each blot (Q: n=4) (c, d) Bioluminescent images of BALB/c mice at 48 hours after treatment of 0.5 mg/kg mRNA delivered through intratracheal instillation of eLNP/Fluc. (c) *In vivo* and (d) *ex vivo* images of luciferase expression. (e) Western blot of hsACE2 protein in the bronchoalveolar lavage fluid (BALF). BALB/c mice were untreated or transfected with *hsACE2* mRNA at 0.75 mg/kg mRNA through intratracheal instillation of eLNP/hsACE2, and the BALF was harvested after 24 h or 48 h post-administration (n = 3). For enrichment of the hsACE2 protein, BALF was subjected to the immunoprecipitation using anti-V5 antibody prior to western blot.

Next, we evaluated whether there was a physical interaction of hsACE2 protein with the RBD of SARS-CoV-2. 293T cells were transfected with eLNP/hsACE2 for 24 h, and untreated cells served as controls. Cell-free conditioned media was collected and inoculated with either PBS or the recombinant His-tagged RBD **(Fig. 4a, b)**. Co-immunoprecipitation was performed with the samples using anti-V5 antibody to capture hsACE2^V5^ **(Fig. 4b, c)**. SDS-PAGE was performed with the immunoprecipitated samples and the flow-through samples, followed by immunoblotting with anti-V5 and anti-His antibodies. It was observed that anti-V5 antibody was able to immunoprecipitate the hsACE2^V5^ from cell-free conditioned media **(Fig. 4c)** while the unbound hsACE2 ^V5^ was detected in the flow-through samples **(Fig. 4d)**. The RBD^His^ was detected only in the immunoprecipitated samples that had both hsACE2 and the RBD **(Fig. 4c)** while the samples that contained RBD^His^ or hsACE2 alone showed no RBD^His^ band in the immunoprecipitated samples **(Fig. 4c)**. The unbound RBD^His^ and hsACE2 were clearly detected in the flow-through samples **(Fig. 4d)**. We also conducted a reciprocal co-immunoprecipitation, in which anti-His antibody was used to capture RBD^His^ **(Fig. 4e, f)**. It was observed that anti-His antibody co-immunoprecipitated hsACE2^V5^ in the samples that contained both hsACE2^V5^ and RBD^His^. hsACE2^V5^ band did not appear in the samples from other groups in immunoblotting. These results demonstrate that hsACE2 protein formed a protein complex with a specific and high-affinity association with the RBD of SARS-CoV-2. We further explored the ability of hsACE2 to inhibit the transduction of the virus using a pseudovirus neutralization assay. To investigate hACE2-dependent infection of the pseudovirus, we created 293T cells stably expressing hACE2 (293T-hACE2) with a lentiviral vector. Expression of hACE2 in the transduced cells was examined in western blot, which showed hACE2 band at approximately 115 kDa (**Extended Data Fig. 7a**). For pseudotyping the SARS-CoV-2, we utilized *Fluc*-packaged HIV-based lentiviral particles containing the S protein of SARS-CoV-2 on them. The lentiviral particles with vesicular stomatitis virus G protein (VSV-G) instead of the S protein were prepared as a positive control. We found the spike pseudovirus infection was hACE2-dependent; while VSV-G pseudovirus infection was not affected by hACE2 expression in host cells (**Extended Data Fig. 7b**). After we examined the hACE2 specificity of the spike pseudovirus, we studied the effects of hsACE2 on the pseudovirus infection. To do this, hsACE2 conditioned media was produced by transfecting *hsACE2* mRNA in 293T/17 cells. The pseudovirus and conditioned media containing hsACE2 were co-incubated on 293T-hACE2 cells and we measured viral transduction. It was found that cells treated with hsACE2 conditioned media led to a drastic reduction (more than 95%) of pseudovirus transduction as compared to the cells treated with pseudovirus alone (**Fig. 4g**). The VSV-G pseudovirus transduction was not affected by either treatment (**Fig. 4h**). These data highlight the potent inhibitory effect of hsACE2 treatment on the SARS-CoV-2 infection by blocking the S protein-mediated viral attachment.

**Fig. 4.**
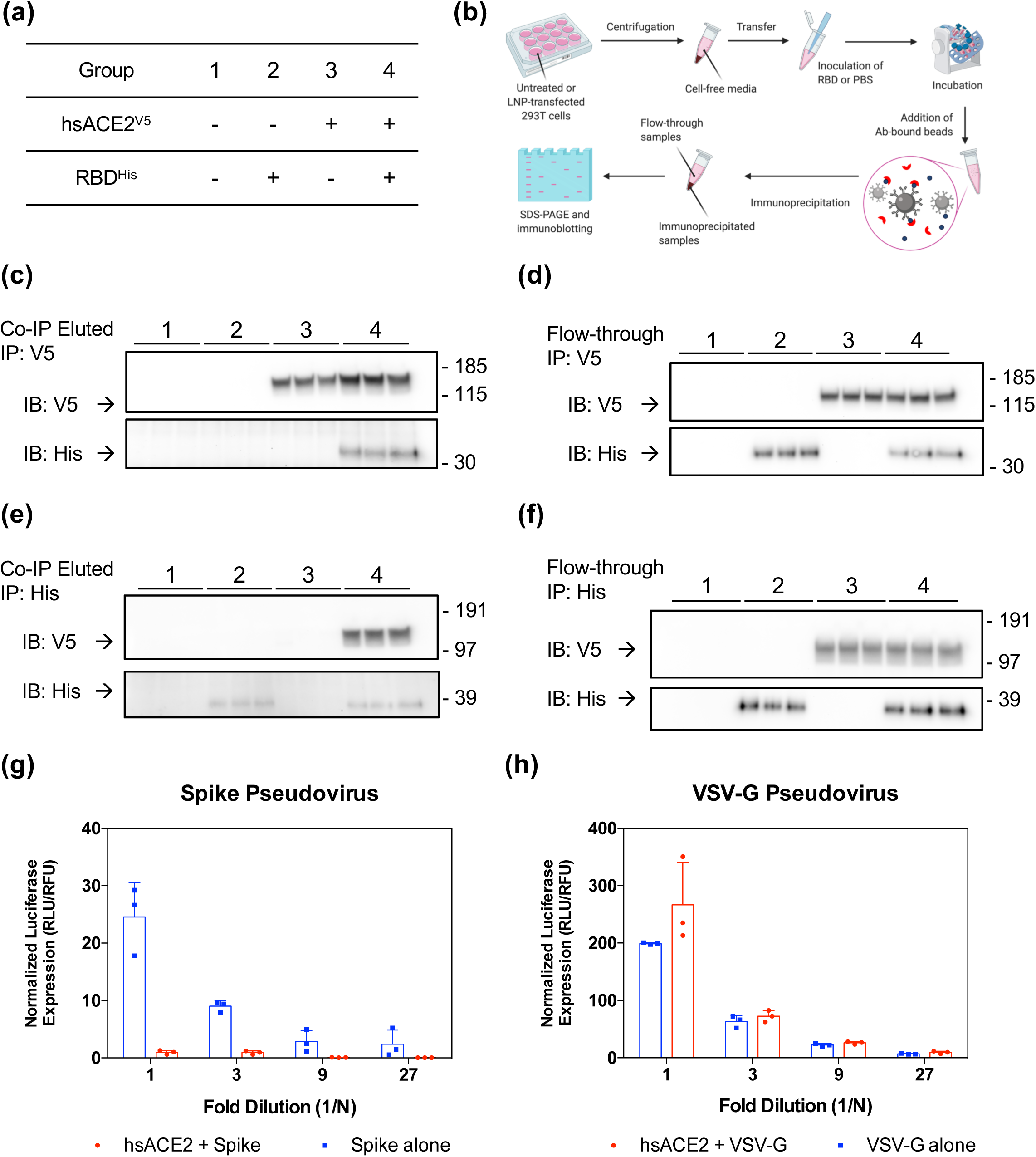
hsACE2 protein binds the RBD and prevents S1-pseudovirus infection. (a) Cell-free media from untreated or hsACE2 transfected 293T cell culture were incubated in the presence or absence of the RBD of the SARS-CoV-2 prior to co-immunoprecipitation (co-IP). + and - define the presence and absence of the treatment, respectively. (b) Schematic workflow of co-immunoprecipitation. (c, d) Co-IP was performed using anti-V5 tag antibody. Upper and lower blots were probed using anti-V5 tag (for hsACE2) and anti-His tag (for RBD) antibodies, respectively. After co-IP, (c) eluted samples and (d) flow-through samples were analyzed using western blot. (e, f) Co-IP using anti-His tag antibody. Upper and lower blots were probed using anti-V5 tag (for hsACE2) and anti-His tag (for RBD) antibodies, respectively. After co-IP, (e) eluted samples and (f) flow-through samples were analyzed in western blot.

There is an urgent need for developing effective treatment options to tackle this pandemic^19,29^. Remdesivir, an investigational antiviral drug, has shown encouraging evidence in improving time of recovery among patients. The overall mortality rate however remains unchanged while conflicting reports have emerged on clinical outcomes^30,31^. Dexamethasone, an anti-inflammatory steroid repurposed for COVID-19, was shown to lower mortality in the patients when used in conjunction with respiratory support^32^. In this study, we demonstrate that mRNA-based nanotherapeutic produces the decoy hsACE2 protein to potentially inhibit the SARS-CoV-2 infection. Using a potent LNP formulation (eLNP), IVT mRNA was delivered to the cytosol and translated into hsACE2 protein more efficiently than the conventional LNPs. hsACE2 protein that was generated from the LNP-delivered mRNA efficiently bound to the RBD of SARS-CoV-2 with a high affinity. Additionally, hsACE2 exerted a potent neutralizing effect on the pseudovirus decorated with the S protein of SARS-CoV-2. Intravenous injection of eLNP/hsACE2 enabled rapid and sustained expression of the circulating hsACE2 protein in the blood circulation within 2 h, which peaked at 6 h and cleared gradually. Lung transfection with eLNP/hsACE2 elicited secretion of hsACE2 protein to the airway mucus in which the primary infection of SARS-CoV-2 occurs. Unlike Fc fragment fused chimeric hsACE2 protein, the availability of mRNA derived circulating hsACE2 is due to continuous generation of new protein from the liver. This provides an opportunity for rapid clearance of the virus while providing protection against the dysregulated RAAS system due to long term presence of newly made protein in the serum. Another use of recombinant hsACE2 was to regulate blood pressure in the Angiotensin II-dependent hypertension^33^. The prevalence of hypertension among the elderly in the United States is more than 60%^34^ and this age-group is also at high risk of COVID-19^35^. In this regard, it is conceivable that expression of enzymatically active hsACE2 from the mRNA therapy could protect COVID-19 patients with hypertension from aggravation of cardiovascular diseases as well as viral infection. Given that binding SARS-CoV-1 to ACE2 results in reduced receptor expression that is associated with SARS-associated ARDS and soluble ACE2 provides protection against virus mediated lung injury^4,36^, it is likely that sustained expression of hsACE2 during infection could facilitate ACE2-mediated lung protection, reduce the incidence of ARDS by neutralizing SARS-CoV-2, and prevent RAAS dysregulation. Additionally, hsACE2 is posited to bind SARS-CoV-2 in the bloodstream and reduce their ability to infect other peripheral organs. It is possible that, by binding and thus masking the RBD, hsACE2 may decrease the amplitude of inflammatory response that causes multiorgan failure.

Our studies are currently limited to the interactions of mRNA-derived hsACE2 with the viral RBD and the ability of hsACE2 mRNA therapy to inhibit cellular entry of the SARS-CoV-2 pseudotyped virus. The affinity of hsACE2 protein to the S protein on the envelop of SARS-CoV-2 could be different, which could have an effect on inhibition of the virus infection. To address these questions, we plan to continue studies that will inform whether LNP-delivered hsACE2 can inhibit live SARS-CoV-2 *in vitro*. For *in vivo* validation, we intend to use hACE2-transgenic animals that express similar pathogenesis when infected with the SAR-CoV-2^37,38^. With the further characterization of mRNA-based hsACE2 treatment, the continued development of inhalable forms of LNPs could enable ease of repeated administration to the patients until the pulmonary infection subsides.

An array of mRNA-based vaccines are progressing with unprecedented speed to effectively end this pandemic^19,20^. mRNA-based nanotherapeutics that produce hsACE2 or neutralizing antibodies in combination with antiviral therapies can emerge as treatment options against SARS-CoV-2 infection. Furthermore, the mRNA-based approach can lead to the development of treatments based on high affinity hsACE2 variants that bind the RBD of SARS-CoV-2 or other hACE2-exploiting coronaviruses that may spill in the future. We hope this work brings to light the potential of mRNA-based therapeutics to treat patients suffering with COVID-19.

## METHODS

### Materials

*Fluc* mRNA and *hsACE2* variant mRNA were purchased from TriLink Biotechnologies (CA, USA). Uridine of *Fluc* mRNA was fully substituted with 5-methoxyuridine, and uridine and cytidine of hsACE2 mRNA were fully substituted with pseudouridine and 5-methyl-cytidine, respectively. Cholesterol and β-sitosterol were purchased from Sigma-Aldrich. DMG-PEG_2K_ was bought from NOF America. DLin-MC3-DMA and DSPC were obtained from BioFine International Inc. and Avanti Polar Lipids, Inc., respectively.

### LNP formulation and characterization

LNPs composed of DLin-MC3-DMA, Cholesterol or β-sitosterol, DMG-PEG_2K_, DSPC, and mRNA were prepared using microfluidic mixing as described previously^39^. Briefly, mRNA was diluted in sterile 50 mM citrate buffer, and lipid components were prepared in 100% ethanol at 50:38.5:1.5:10 molar ratio. The lipid and mRNA solutions were mixed using the NanoAssemblr Benchtop at a 1:3 ratio, followed by overnight dialysis against sterile PBS using a Slide-A-Lyzer G2 cassette with 10,000 Da molecular-weight-cut-off (Thermo Fisher Scientific). Dialyzed LNP solutions were concentrated using Amicon^®^ Ultra centrifugal filter units with 10,000 Da molecular-weight-cut-off (Millipore). Hydrodynamic size and PDI of the LNPs were measured in dynamic light scattering using the Zetasizer Nano ZSP (Malvern Instruments, UK). mRNA encapsulation was assayed using a Quant-iT™ RiboGreen^®^ RNA Assay kit (Thermo Fisher Scientific) and a multimode microplate reader (Tecan Trading AG, Switzerland).

### Cell culture

293T, Calu-3, Hep G2 cell lines were kindly gifted from Prof. Sadik Esener (OHSU), Prof. Kelvin MacDonald (OHSU), and Prof. Conroy Sun (OSU), respectively. 293T/17 cell line was purchased from ATCC (CRL-11268). 293T, 293T/17 and Hep G2 cells were cultured in DMEM supplemented with 10% heat-inactivated FBS and 1% penicillin/streptomycin. Calu-3 cells were cultured in MEM supplemented with 10% heat-inactivated FBS, 1% penicillin/streptomycin, non-essential amino acids, and sodium pyruvate.

### *In vitro Fluc* mRNA transfection assay

For *in vitro Fluc* mRNA transfection assays, cells were seeded on a white 96 well plate at 4×10^3^ cells/well for 293T and Hep G2 cells or at 10^4^ cells/well for Calu-3, followed by overnight incubation for cell attachment. Cells were incubated with nanoparticles encapsulating *Fluc* mRNA and analyzed for cell viability and luciferase activity with the ONE-Glo™+Tox luciferase reporter and cell viability assay kit (Promega) using a multimode microplate reader.

### *In vitro hsACE2* mRNA transfection

For *in vitro* mRNA transfection for hsACE2 production, cells were seeded on a 12-well plate at 3×10^5^ cells per well and allowed to attach for overnight. Cells were treated with LNPs encapsulating hsACE2 mRNA for 24 h, and culture media were centrifuged at 500 *g* for 10 min at 4°C. Cell-free media was supplemented with protease and phosphatase inhibitor cocktail (Thermo Fisher Scientific) and used for downstream experiments.

Besides culture media, transfected cells were lysed using RIPA buffer containing protease and phosphatase inhibitor cocktail, followed by centrifugation at 16,000 *g* for 30 min at 4°C. Supernatant lysate was collected for western blot.

### Detection of hsACE2 protein by western blot

Production of hsACE2 protein upon transfection was detected by western blot. In brief, total protein concentration of sample was quantified using a Micro BCA protein assay kit (Thermo Fisher Scientific) according to the manufacturer’s instruction. Cell-free supernatants or cell lysates containing 30 µg of total protein were prepared in 1X LDS sample buffer under reducing conditions, denatured at 70°C for 10 min, and run on 4- 12% Bis-Tris gels or 4-20% Tris-glycine gels, followed by dry transfer to PVDF membrane using iBlot 2 Dry Blotting System (Thermo Fisher Scientific). The blots were blocked using 5% skim milk for 1 h at room temperature. The primary antibodies used were: rabbit monoclonal anti-V5 tag at 1:1,000 (Cell Signaling Technology, 13202), rabbit monoclonal anti-6x-His tag at 1:1,000 (Thermo Fisher Scientific, MA5-33032), and mouse monoclonal anti-β-actin at 1:10,000 (R&D Systems, MAB8929). The secondary antibodies used were goat polyclonal anti-rabbit HRP (Jackson ImmunoResearch, 111-035-003) and anti-mouse HRP (115-035-003). For detection and documentation, we used SuperSignal™ West Pico Plus Chemiluminescent Substrate and myECL imager (Thermo Fisher Scientific). After chemiluminescent imaging, blots were further stained using GelCode™ Blue Safe Protein Stain (Thermo Fisher Scientific) according to the manufacturer’s instruction.

### Co-immunoprecipitation of hsACE2 and SARS-CoV-2 Spike RBD

Cell free media from untreated or transfected 293T cell culture was prepared. 1 µg of SARS-CoV-2 Spike RBD-His (Sino Biological) was inoculated to 400 µl of cell free media, followed by overnight incubation at 4°C with rotation. Subsequent co-immunoprecipitation was conducted using Dynabeads™ Protein G Immunoprecipitation kit (Thermo Fisher Scientific) according to the manufacturer’s instruction. Briefly, cell-free media inoculated with the spike RBD were incubated with antibody bound Dynabeads for 20 min at room temperature with rotation. The antibodies used for pull-down were mouse monoclonal anti-His tag (sc-8036) or anti-V5 tag (sc-81594) antibody (Santa Cruz Biotechnology). Following three washes with PBS, samples were eluted using elution buffer and denatured using LDS sample buffer and reducing agent for western blot.

### Animals

All animal studies were conducted at Oregon Health and Sciences University and approved by the Institutional Animal Care and Use Committee (IACUC, IP00001707).

### *In vivo Fluc* mRNA transfection via intravenous administration

Female BALB/c mice (8-12 weeks) were sedated using isoflurane, and LNPs encapsulating *Fluc* mRNA were intravenously administered via tail vein. At predetermined time points post-administration, 200 µl of D-luciferin substrate was intraperitoneally injected to the mice 10 minutes prior to bioluminescence imaging (150 mg/kg). Image acquisition and analysis were performed using the IVIS^®^ Lumina XRMS and the manufacturer’s software (PerkinElmer).

### *In vivo hsACE2* mRNA transfection via intravenous administration

Female BALB/c mice (8-12 weeks) were sedated using isoflurane, and LNPs encapsulating *hsACE2* mRNA were administered to animals via tail vein. At predetermined time points post-administration, whole blood was collected using cardiac puncture or submandibular bleeding. The collected blood samples were processed to sera using serum-separating tubes (BD). The separated sera were used for downstream experiments. Mouse liver were sterilely harvested and homogenized using a handheld tissue homogenizer.

### *In vivo* mRNA transfection via intratracheal instillation

Intratracheal instillation was performed as described previously^40^. Female BALB/c mice (8-12 weeks) were anesthetized using ketamine/xylazine cocktail. Anesthetized animals were leaned over intubation stand (Kent Scientific), and their vocal cords were directly visualized using an otoscope with a 2-mm speculum (Welch Allyn). A flexible guide wire was advanced through the vocal cords to trachea. Once the wire was located within trachea, a 20 G catheter was passed over the wire and the wire was removed. To administer LNPs, a gas tight syringe with a 22 G blunt needle (Hamilton) was filled with LNPs containing mRNA. The syringe were inserted through the catheter and LNPs encapsulating mRNA was administered to lungs, followed by 100 µl of air to distribute the LNP solution throughout the lungs.

### Collection of bronchoalveolar lavage fluid (BALF)

After intratracheal instillation, euthanize the animals by CO_2_ asphyxia at an appropriate time post-administration. The trachea was surgically exposed and intubated with a 20 G catheter. The mouse lungs lavage was performed three times with 0.8 ml of prewarm PBS to collect BALF. The collected BALF was centrifuged at 500 *g* for 10 min at 4°C. The supernatants were supplemented with protease and phosphatase inhibitor cocktail and used for downstream experiments.

### Immunoprecipitation of hsACE2 from BALF

Immunoprecipitation of hsACE2 from BALF was conducted using Dynabeads™ Protein G Immunoprecipitation kit (Thermo Fisher Scientific) according to the manufacturer’s instruction. The collected BALF was incubated with Dynabeads having anti-V5 tag antibody for 20 min at room temperature with rotation. Following three washes with PBS, samples were eluted using elution buffer and denatured using LDS sample buffer and reducing agent for western blot.

### Plasmids

Lentiviral reporter plasmid pHAGE-CMV-Luc2-IRES-ZsGreen-W (BEI Resources, NR-52516) and helper plasmids pHDM-Hgpm2 (BEI Resources, NR-52517), pHDM-tatb (BEI Resources, NR-52518), pRC-CMV-rev1b (BEI Resources, NR-52519), and hACE2 containing pHAGE2-EF1a ACE2 (BEI Resources, NR-52512) were kindly provided by Jesse D. Bloom^41^. pcDNA3.1-SARS2-Spike (Addgene, #145032) was a gift from Fang Li^42^. pMD2.G containing VSV-G envelop protein (Addgene, #12259) and pCMVΔR8.2 (Addgene, #12263) were a gift from Didier Trono.

### Generation of 293T-hACE2

In order to create 293T/17 cells overexpressing hACE2, we transduced the hACE2 gene to 293T/17 cells using a lentiviral vector. To produce the lentivirus packaging hACE2 gene, 293T/17 cells were transfected with pCMVΔR8.2, pMD2.G, and pHAGE2-EF1aInt-ACE2-WT using lipofectamine 2000. After 4 h, the cells were replenished with the fresh growth media. After 48 h, the lentiviral particles were collected, filtered, and used immediately to transduce 293T/17 cells. After 48 h transduction, the cells were harvested, passaged with the growth media, and referred as 293T-hACE2. To confirm the expression of hACE2 after transduction, 293T-hACE2 cells were harvested and lysed using RIPA buffer containing protease and phosphatase inhibitor cocktail. Cell lysates was processed to perform western blot analysis as described above. To probe hACE2, anti-ACE2 antibody (Santa Cruz Biotechnology, sc-390851) and anti-mouse HRP were used as the primary and secondary antibodies at 1:200 and 1:2,000, respectively.

### Production of pseudovirus particles

293T/17 cells were seeded in T-75 flask at 5×10^4^ cells/flask and grown for 18 h. Cells were co-transfected with 7.8 µg of the lentiviral reporter, 1.7 µg of each helper plasmids, and either 7.8 µg of pcDNA3.1-SARS2-Spike (Spike), 2.5 µg of pMD2.G (VSV-G), or no plasmid (No envelope) using lipofectamine 3000 as instructed by manufacture. After 48 h, pseudoviruses were collected, filtered, aliquoted into single-use vials, and stored at - 80°C.

### Titration of pseudovirus particles

293T-hACE2 cells were seeded at 10^4^ cells/well in white, 96-well plates and grown for 18 h. Cells were transduced in triplicate with a 4-point, 1:3 serial dilution of the pseudoviruses with polybrene at a final concentration of 5 µg/ml. Polybrene was not included in the VSV-G pseudovirus-treated wells. After 48 h, cell viability and luciferase activity were assessed with the ONE-Glo™+Tox luciferase reporter and cell viability assay kit.

### Preparation of conditioned media for pseudovirus neutralization assay

To make conditioned media containing hsACE2, 293T/17 cells were seeded into T-75 flasks at 5×10^4^ cells/flask and grown 18 h. Cells were transfected with 22 µg mRNA or equivalent volume of PBS using lipofectamine 3000. After incubation for 6 h, cells were washed with PBS and the complete media was added. After 24 h, media was harvested, filtered with 0.45 um filter, and concentrated in a spin column with Amicon^®^ Ultra centrifugal filter units with 10,000 Da molecular-weight-cut-off at 4,000 *g* for 30 minutes. The concentrated, conditioned media was brought up to 2 ml with serum-free media and used immediately in the neutralization assay.

### Pseudovirus neutralization assay

For neutralization assay, 293T-hACE2 cells were seeded into white 96-well plates at 2×10^4^ cells/well and grown for 24 h. Pseudovirus was serially diluted as before. The conditioned media was added to the serial dilutions at ratio of 2 : 3 for conditioned media : pseudovirus, and incubated at 4°C for 1 h. Polybrene was added as before. Media was removed from the 96-well plates and cells were transduced as before. After 48 h, cell viability and luciferase activity were assessed with the ONE-Glo™+Tox luciferase reporter and cell viability assay kit.

## ACKNOWLEDGEMENTS

This project was supported through funding from OSU College of Pharmacy startup funding (G.S). Sahay lab receives support through the National Heart Lung and Blood Institute (N.H.L.B.I) 1R01HL146736-01 (G.S) and Cystic Fibrosis Foundation - SAHAY18G0 (G.S) and SAHAY 19XX0. We thank Dr. Jesse Bloom at Fred Hutchinson Cancer Research Center for kindly providing the plasmids for the pseudovirus neutralization assay. We also thank Dr. Jessica Smith and Dr. Alec Hirsch from the Vaccine and Gene Therapy Institute for feedback.

## AUTHOR CONTRIBUTIONS

G.S. directed this research. G.S. and A.M conceived the idea. G.S, A.M and J.K. designed the experiment. J.K performed all experiments except the pseudovirus transduction with feedback from A.M and G.S. D.N and A.J made the pseudovirus with feedback from G.S. and performed inhibition assays. J.K made all the figures with feedback from all authors. G.S, J.K and A.M wrote the paper with feedback from all authors.

## COMPETING INTERESTS

G.S. is a co-inventor in patent application US20200129445A1 that details eLNPs.

---

**Extended Data Fig. 1:**
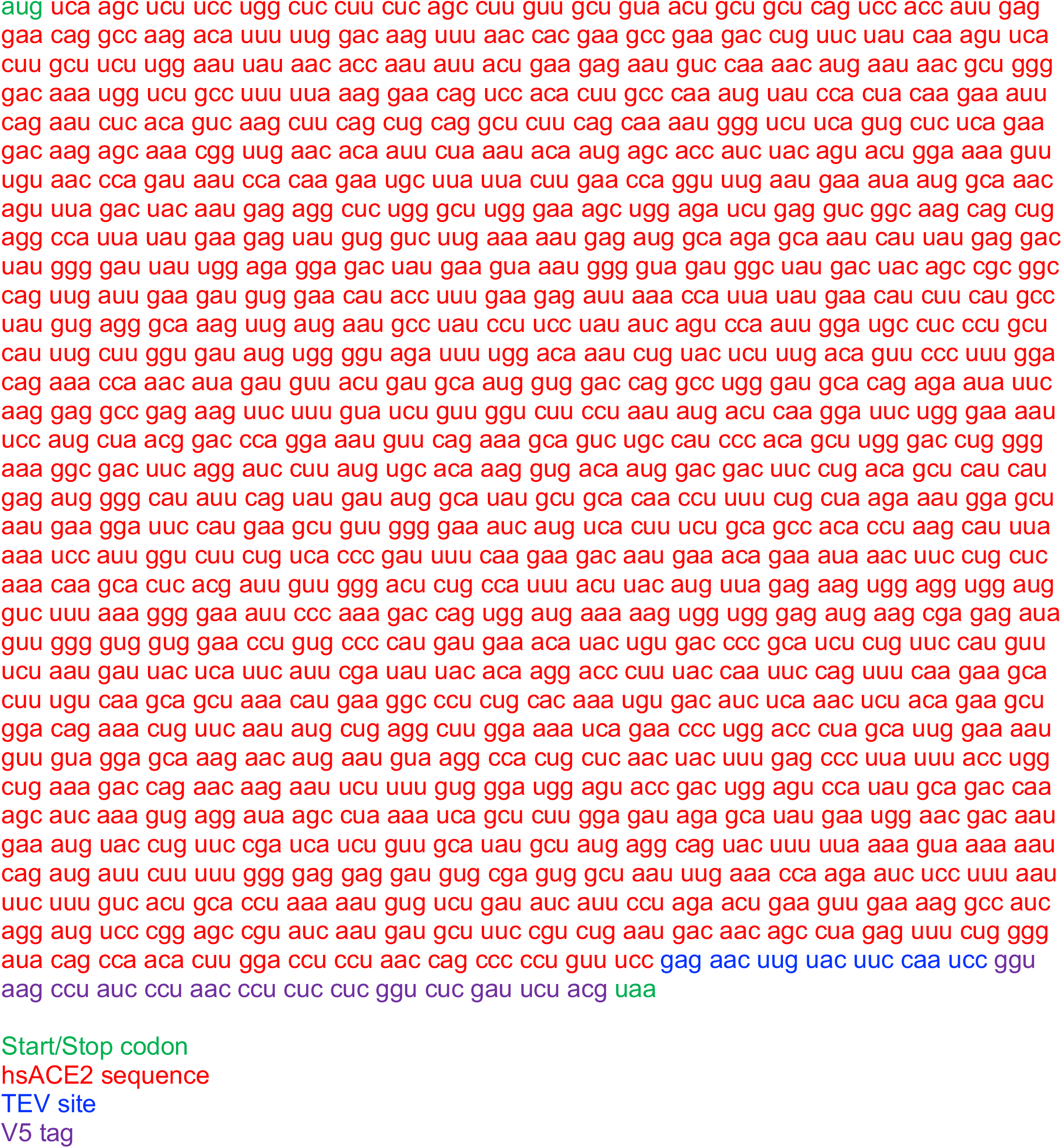
mRNA sequence of hsACE2 variant. Green bases are start and stop codons. Red bases encodes the 1-740 amino acid sequence of human ACE2 protein. Blue bases encodes TEV protease site that can be used as a cleavage site for the removal of the C-terminal tag. Purple bases encodes the V5 epitope tag.

**Extended Data Fig. 2:**
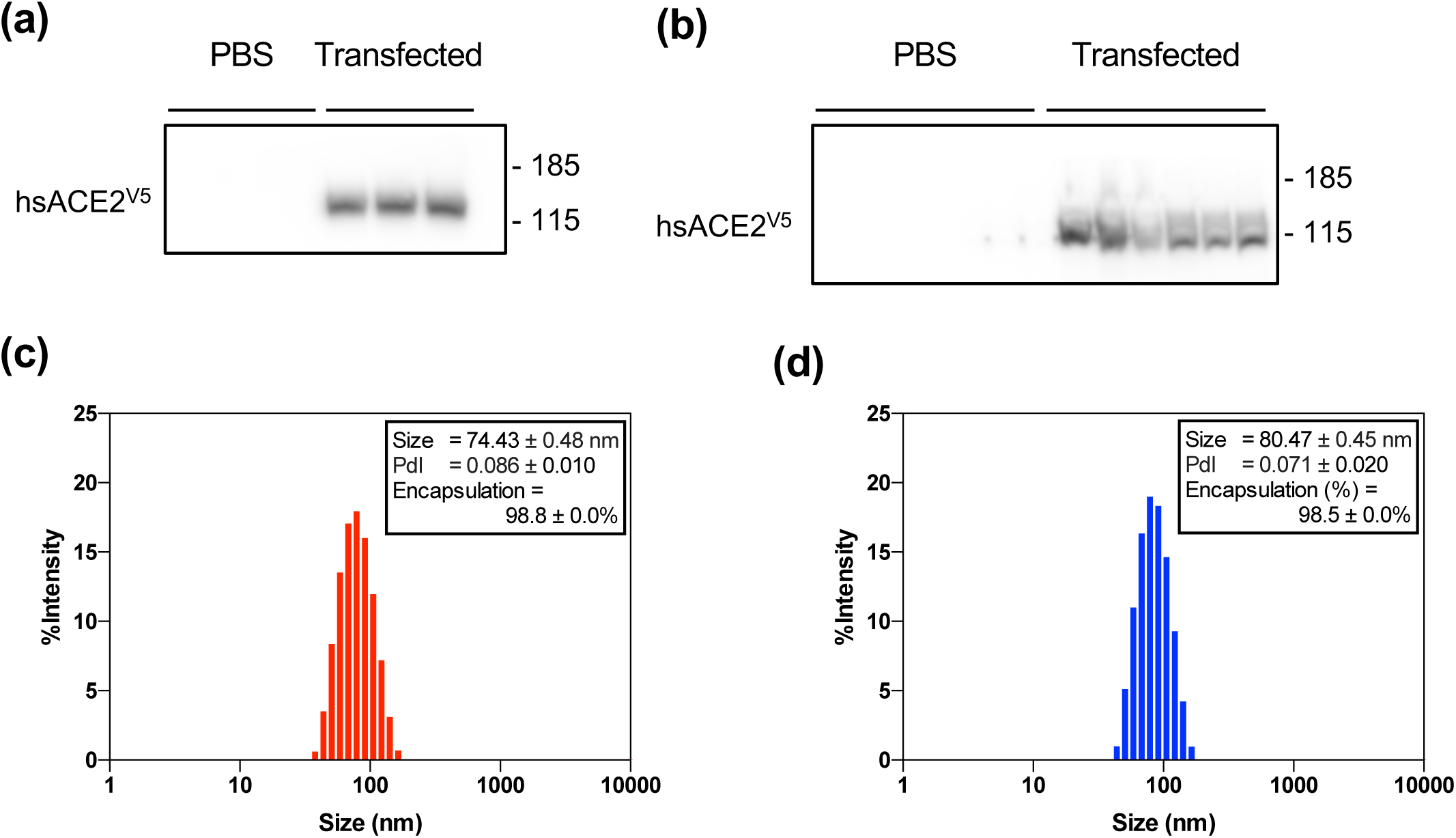
mRNA derived hsACE2 protein expression and characterization of LNPs encapsulating the mRNA. (a, b) Western blot of (a) cell-free media and (b) cell lysates derived from 293T cell culture that was transfected with hsACE2 mRNA using lipofectamine 3000 for 24 hours. (c, d) Representative data of size distribution and RNA encapsulation of (c) LNP/hsACE2 and (d) eLNP/hsACE2 that were used in the study.

**Extended Data Fig. 3:**
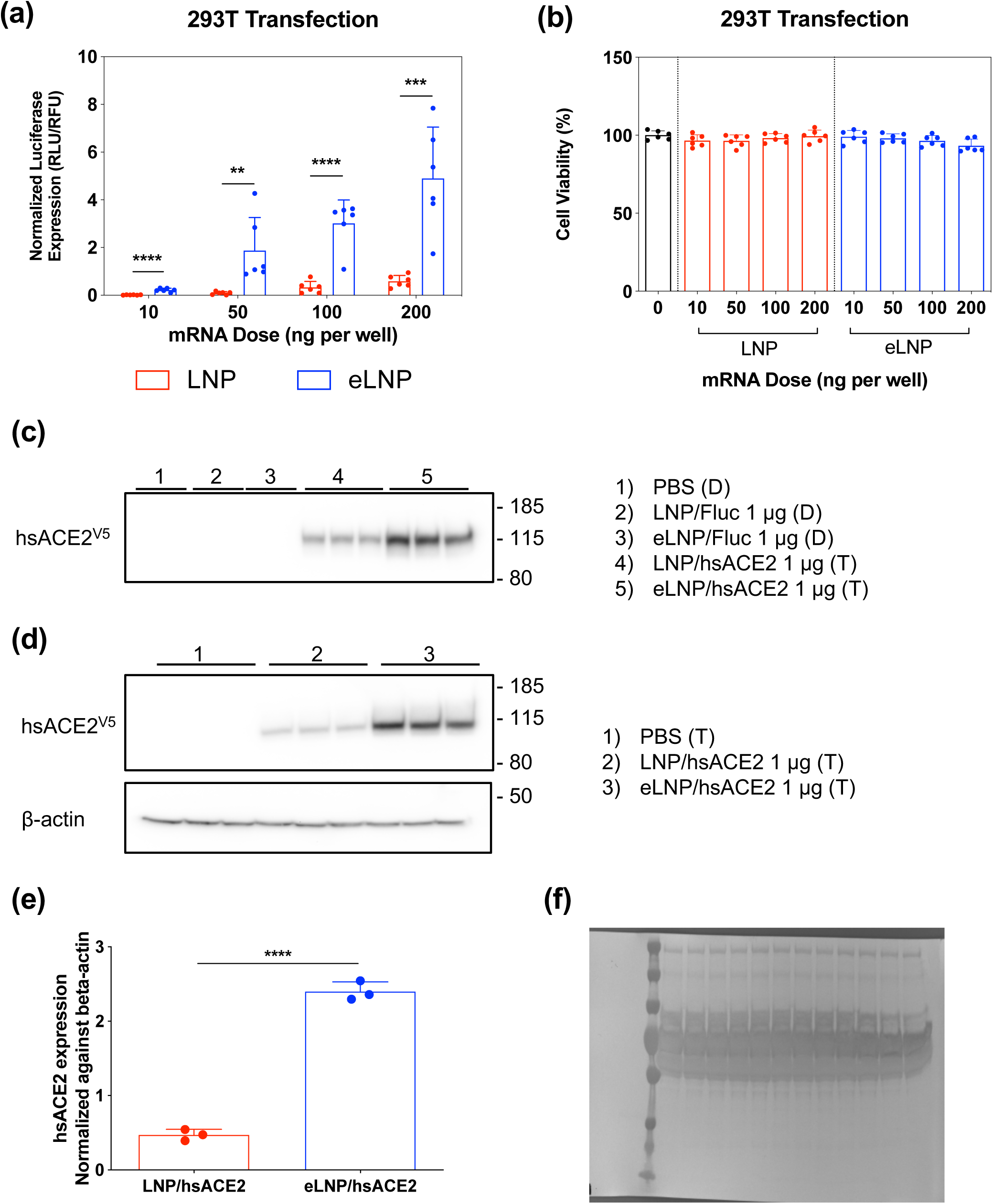
Nanoparticle-delivered mRNA transfection in 293T cells. (a) *In vitro* luciferase and (b) cell viability assay of 293T cells transfected with LNP/Fluc or eLNP/Fluc for 24 h (10-200 ng mRNA per well, n = 6). (c) Western blot with cell-free media and (d) cell lysates of 293T cells after mRNA transfection using various LNPs. Treatment and mRNA dose are described on the right of each blot (e) Expression of hsACE2 protein in the 293T cell lysates was normalized to the expression of β-actin by densitometry. (f) The blot used to probe the Fig. 2a was stained with Coomassie blue to visualize total protein. Statistical analysis was performed using Student’s *t* test. \*\**p* < 0.01, \*\*\**p* < 0.001, \*\*\*\**p* < 0.0001. All data were expressed as the mean ± S.D.

**Extended Data Fig. 4:**
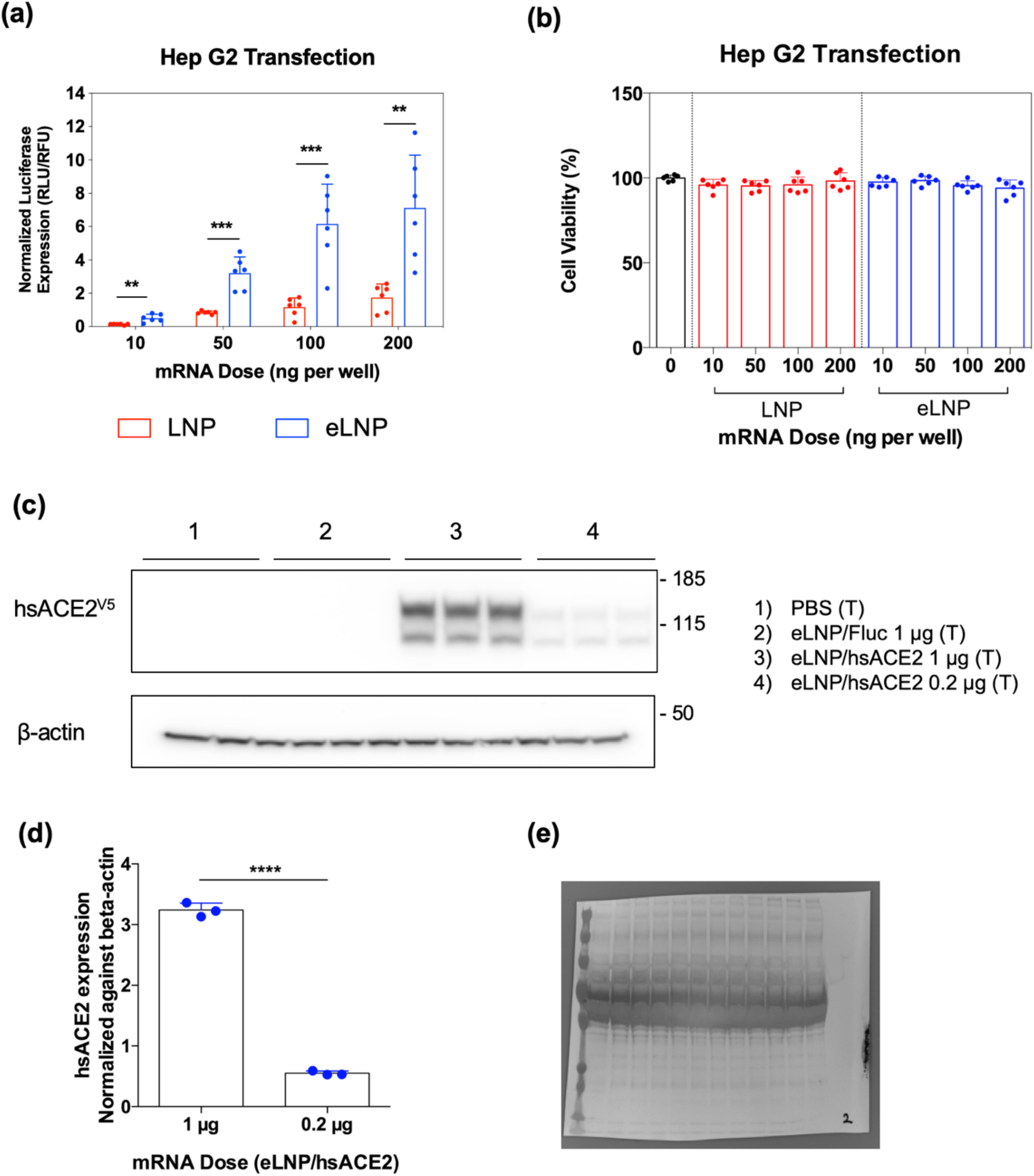
Nanoparticle-delivered mRNA transfection in Hep G2 cells. (a) *In vitro* luciferase and (b) cell viability assay of Hep G2 cells transfected with LNP/Fluc or eLNP/Fluc for 24 h (10-200 ng mRNA per well, n = 6). (c) Western blot with cell lysates of Hep G2 cells after mRNA transfection using various LNPs. Treatment and mRNA dose are described on the right of each blot (T; n=3). (d) Expression of hsACE2 protein in the Hep G2 cell lysates was normalized to the expression of β-actin by densitometry. All data were expressed as the mean ± S.D. (e) The blot used to probe the Fig. 2b was stained with coomassie blue to visualize total protein. Statistical analysis was performed using Student’s *t* test. \*\**p* < 0.01, \*\*\**p* < 0.001, \*\*\*\**p* < 0.0001. All data were expressed as the mean ± S.D.

**Extended Data Fig. 5:**
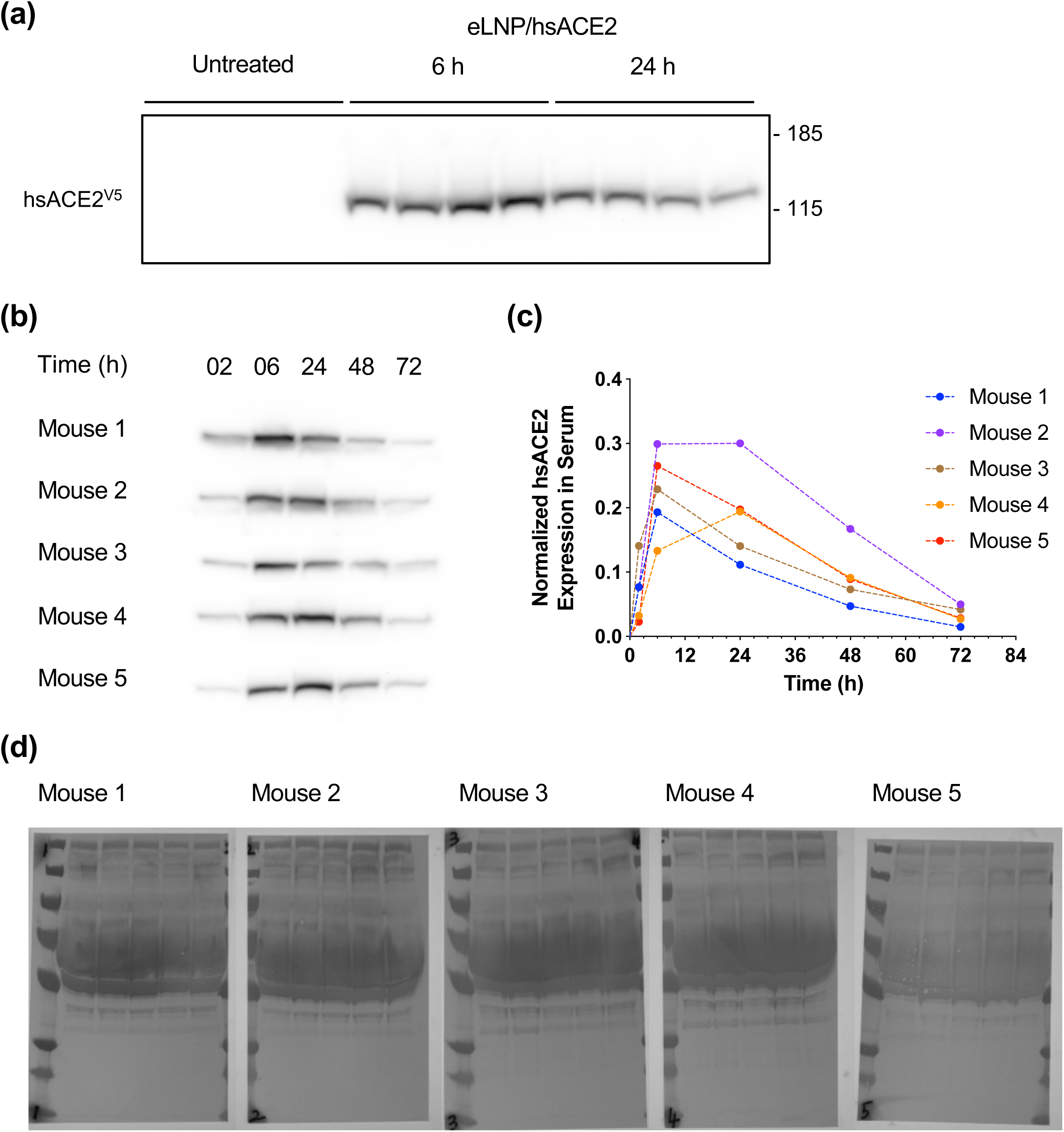
*In vivo* expression of circulating hsACE2 protein post-intravenous administration of eLNP/hsACE2. (a) Western blot of hsACE2 protein in mouse sera after IV injection of eLNP/hsACE2. (b) Western blot with mouse sera collected with predetermined time intervals after IV injection of eLNP/hsACE2 (n = 5). (c) Temporal level of the circulatory hsACE2 protein in each mouse. Expression of the hsACE2 protein was normalized to the total amount of protein in each lane. (d) The blots used to probe the Extended Data Fig. 5b were stained with Coomassie blue to visualize total protein.

**Extended Data Fig. 6:**
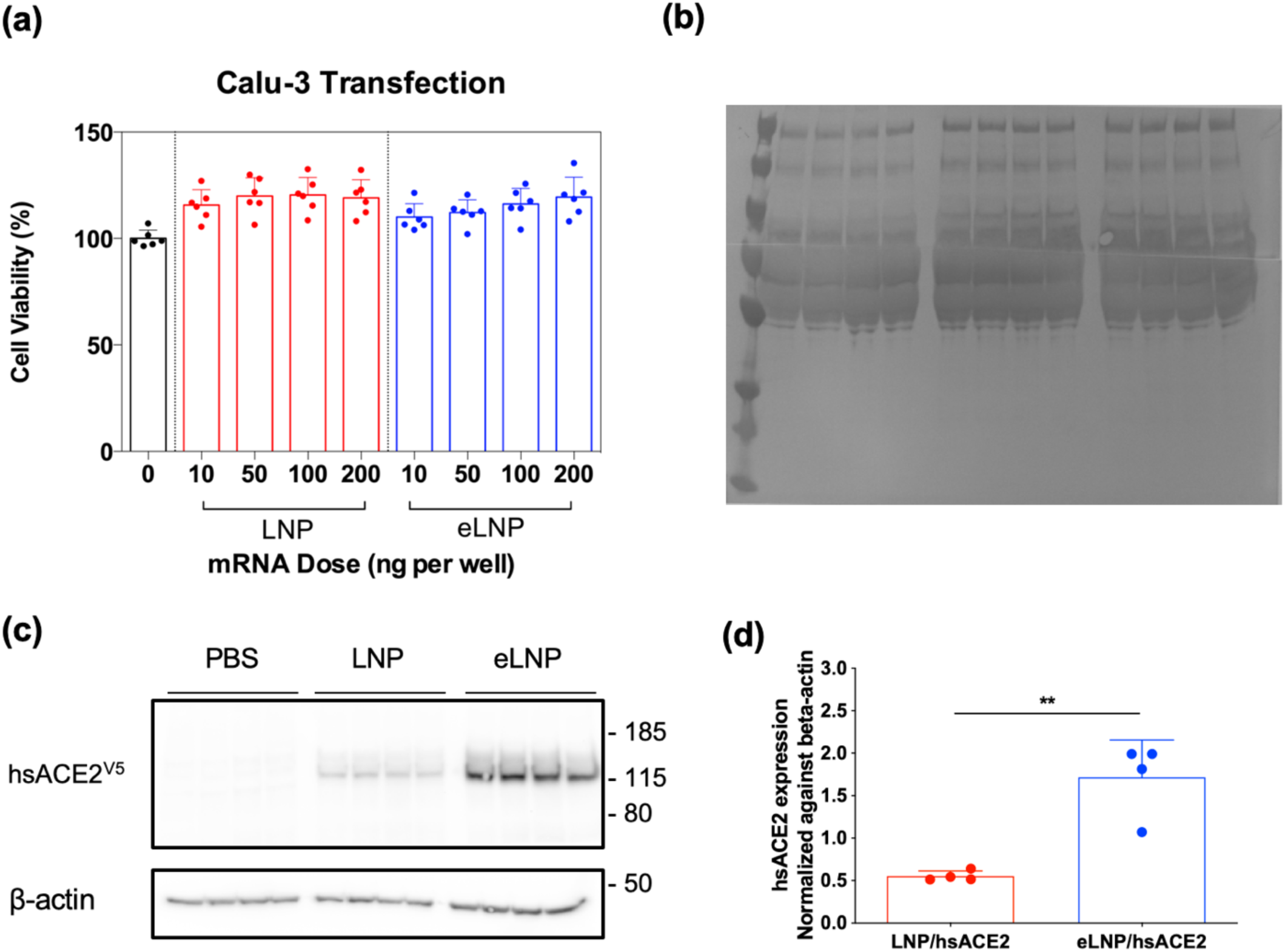
LNP-delivered mRNA transfection in Calu-3 cells. (a) Cell viability of Calu-3 cells transfected with LNP/Fluc or eLNP/Fluc for 48 h (10-200 ng mRNA per well, n = 6). (b) The blots used to probe the Fig. 4b were stained with Coomassie blue to visualize total protein. (c) Western blot with cell lysates of Calu-3 cells after mRNA transfection using LNP/hsACE2 and eLNP/hsACE2 (n = 4). (d) Expression of hsACE2 protein in the Calu-3 cell lysates was normalized to the expression of β-actin by densitometry. Statistical analysis was performed using Student’s *t* test. \*\**p* < 0.01. All data were expressed as the mean ± S.D.

**Extended Data Fig. 7:**
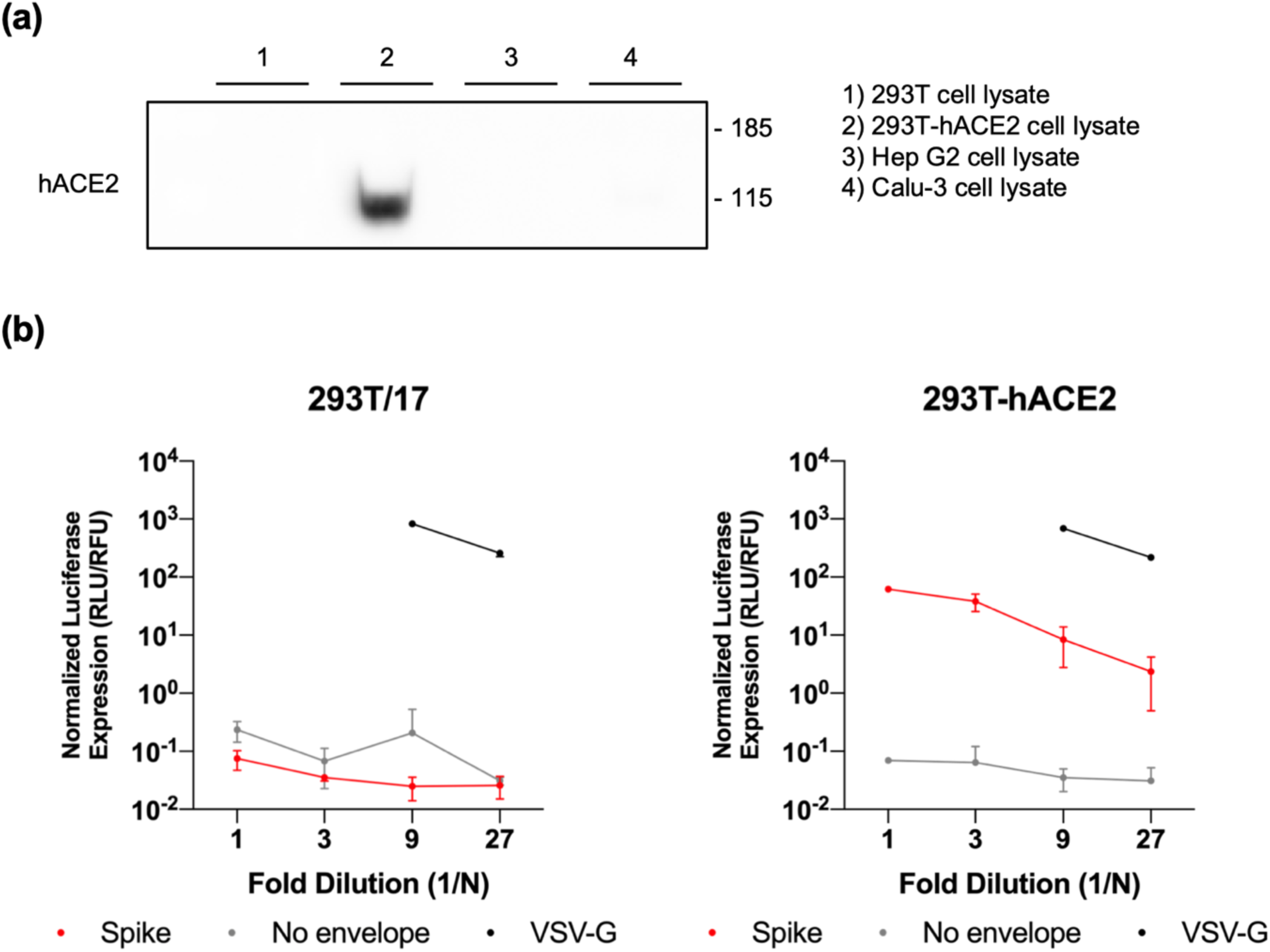
Generation of 293T-hACE2 cells and effects of hACE2 in transduction of the spike pseudovirus. (a) Western blot of hACE2 in various cell lysates. (b) Titration of various pseudovirus in (left) 293T cells and (right) 293T-hACE2 cells (n = 3). Red; spike pseudovirus, gray; pseudovirus with no envelope, black; VSV-G pseudovirus. Normalized luciferase expressions of VSV-G pseudovirus at 1 and 3-fold-dilutions were not presented due to saturation of signal. All data were expressed as the mean ± S.D.

